# Maximized field-of-view deep-brain calcium imaging through gradient-index lenses

**DOI:** 10.1101/2025.09.08.674926

**Authors:** Chenmao Wang, Zongyue Cheng, Yuting Li, Jianian Lin, Meng Cui

**Affiliations:** School of Electrical and Computer Engineering, Purdue University, West Lafayette, IN 47907, USA; Bindley Bioscience Center, Purdue University, West Lafayette, IN 47907, USA; Department of Biology, Purdue University, West Lafayette, IN 47907, USA

## Abstract

Advances in genetically encoded fluorescent indicators have enabled increasingly sensitive optical recordings of neural activity. However, light scattering in the mammalian brain tissue restricts optical access to deeper regions. To address this limitation, researchers often employ implanted gradient-index (GRIN) lenses to reach deep brain areas. Nevertheless, the severe optical aberrations of GRIN lenses significantly reduce the effective field of view (FOV). In this work, we present a simple and robust imaging approach that combines low-NA telecentric scanning (LNTS) of laser excitation with high-NA fluorescence collection to increase the FOV. This configuration effectively eliminates common aberrations such as astigmatism and field curvature, resulting in a FOV ∼100% as large as the GRIN lens facet area — corresponding to a ∼400% increase in imaging area compared with conventional approaches. We validate this method through both structural and functional *in vivo* imaging. The highly consistent imaging performance, fully maximized imaging FOV, and the very simple optical design make this method well-suited for broad dissemination in neuroscience research.

## Introduction

Observing neuronal activity at high spatial and temporal resolutions in the brains of behaving animals is essential for neuroscience research. The rapid development of genetically encoded fluorescent indicators^1-3^ has enabled the widespread adoption of optical imaging techniques,^4-7^ allowing researchers to monitor activity with cellular to subcellular resolution. However, a major challenge of this approach is the limited imaging depth in mammalian brains, primarily due to light scattering in biological tissue^8^. Even with multiphoton excitation, *in vivo* calcium imaging typically penetrates approximately one millimeter^9, 10^, leaving most brain regions inaccessible. To address this limitation, miniature optical implants have emerged as a common strategy for accessing deeper brain structures. Among these, gradient-index (GRIN) lenses are widely favored due to their compact design and ease of use.^11-28^ In particular, GRIN lenses with a diameter of 0.5 mm and a length of approximately 7 mm are commonly used for deep brain imaging in mice.

A fundamental challenge of using GRIN lenses for deep-brain imaging is the limited imaging FOV. GRIN lenses inherently suffer from significant fourth-order astigmatism^29^, characterized by the term W_222_(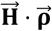)^2^, where W_222_ represents the fourth-order aberration coefficient (i.e., the magnitude of the aberration),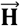 represents the normalized field position and 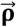 represents the normalized pupil position. Because the aberration scales quadratically with field position, its severity increases toward the edges of the FOV, degrading spatial resolution, signal intensity, and image contrast.^30^ This fourth-order astigmatism results in three distinct focal planes—sagittal, medial, and tangential—leading to out-of-focus artifacts and background signal. Due to its spatially varying nature, this aberration is difficult to correct using conventional methods, especially during high-speed functional imaging. Consequently, in typical one-photon wide-field imaging, the usable FOV with soma-resolution is confined to roughly half of the lens diameter, amounting to approximately 25% of the lens facet area. When neurons of interest are outside this limited region, additional animals and surgical procedures would be required. Ultimately, the small effective FOV imposes significant constraints on imaging throughput and experimental efficiency of deep brain neuroscience studies.

Several solutions have been explored to expand the usable FOV for imaging through GRIN lenses. One widely investigated approach is adaptive optics (AO)^31-33^, which corrects for aberrations introduced by the GRIN lens. However, due to the spatial variation of the aberration, each correction is only effective within a limited portion of the FOV. As a result, numerous sequential corrections and images are required to cover the entire FOV, making this approach impractical for high-speed functional imaging. Geometric transform adaptive optics (GTAO)^30^ has been developed to address this limitation by enabling continuous, spatially varying aberration correction. It leverages spatial transformations to employ the astigmatism of one GRIN lens to compensate for that of another. Although GTAO can offer spatially varying aberration correction and is compatible with a wide range of GRIN lenses and wavelengths, its optical and mechanical implementation is technically complex. In comparison, lens design-based correction^34^ offers a more practical and scalable solution for GRIN lens imaging. After all, the fourth-order astigmatism and its associated field curvature are within the common vocabulary of lens design. Although the lens design-based solution improves the FOV, the achievable imaging FOV is typically still below 50% of the lens facet area. Differences in lens diameter, pitch length, and pitch number necessitate custom-designed optics for each configuration, reducing flexibility and limiting broad applicability.

To address these challenges and enhance the success rate and throughput of GRIN lens-based deep brain calcium imaging, we present a simple, robust, and adaptable solution capable of providing a FOV ∼ 100% as large as the GRIN lens facet, thereby maximizing the usable FOV to its physical limit.

## Results

### Optical configuration for 100% FOV calcium imaging

Although two-photon imaging offers superior contrast and penetration depth, one-photon calcium imaging through GRIN lenses remains the most widely used solution for its simplicity, lower cost, and compatibility with both head-fixed and freely moving animal studies. For these reasons, we developed a simple and robust solution for one-photon calcium imaging through GRIN lenses, which is adaptable to various GRIN lenses and wavelengths. Similar to conventional optics, the aberrations of the GRIN lens strongly depend on the numerical aperture (NA). A straightforward strategy to reduce aberrations and expand the imaging FOV is to lower the excitation NA (Supplementary Fig. 1). For soma-resolution calcium imaging, reduced spatial resolution is often acceptable, and the extended focal depth can increase the number of visible cells within a single image plane for simultaneous recordings. However, reducing NA can substantially decrease fluorescence signal-collection efficiency and increase susceptibility to photobleaching. For instance, reducing the imaging NA from 0.4 to 0.04 significantly suppresses astigmatism (about 100-fold reduction) but also reduces the signal collection efficiency by a factor of 102. While light-field detection could, in principle, boost the total signal collection by using an array of low-NA apertures, the radially varying refractive index of GRIN lenses prevents high-NA, non-telecentric rays from reaching the periphery of the lens facet, resulting in vignetting. With these factors considered, we proposed to use low-NA telecentric scanning (LNTS) for the excitation with ∼100% FOV coverage and utilized the GRIN lens as a light pipe to collect most of the fluorescence emission with an NA of ∼ 1.0. Such a simple laser scanning configuration has the following properties. First, the low-NA telecentric beam configuration can effectively utilize ∼ 100% of the lens facet for imaging at the maximized FOV coverage without vignetting. Second, the beam encounters negligible aberration, free from the commonly encountered astigmatism and its associated field curvature. Third, the light pipe-based signal collection can effectively utilize the fluorescence emission including the randomly scattered and highly aberrated emission light which would contribute to background in the conventional wide-field imaging. Fourth, the extended focal depth allows the simultaneous calcium recording from more cells.

The optical implementation of this solution is straightforward (Fig. 1a, b). The system consists of a laser-scanning setup using a low-NA (∼0.04) telecentric excitation beam at 488 nm, combined with high-NA (1.0) fluorescence collection across the entire aperture of the GRIN lens. To evaluate the imaging system, we embedded 4 µm fluorescent beads in agar and inserted a GRIN lens of 0.5 mm in diameter and 6.7 mm in length into the sample. Under conventional wide-field recording at an NA of 0.4, beads located near the center of the FOV were well resolved. In comparison, beads positioned at approximately 50% of the lens diameter exhibited pronounced astigmatism, and image quality degraded significantly toward the periphery (Fig. 1c). In comparison, the LNTS approach produced clear images of beads across the entire 500 µm FOV, matching the diameter of the GRIN lens (Fig. 1d). The extended focal depth enabled simultaneous visualization of more beads in the same imaging plane (Fig. 1e) and provided greater tolerance to axial motion and alignment errors. For comparison, we also varied the detection NA from 0.05 to 0.4 for the conventional camera-based wide-field imaging system (Supplementary Fig. 2). As expected, reducing the detection NA to 0.05 allowed imaging closer to the edge of the lens, but at the cost of substantially reduced fluorescence collection efficiency and increased illumination power.

**Figure 1.**
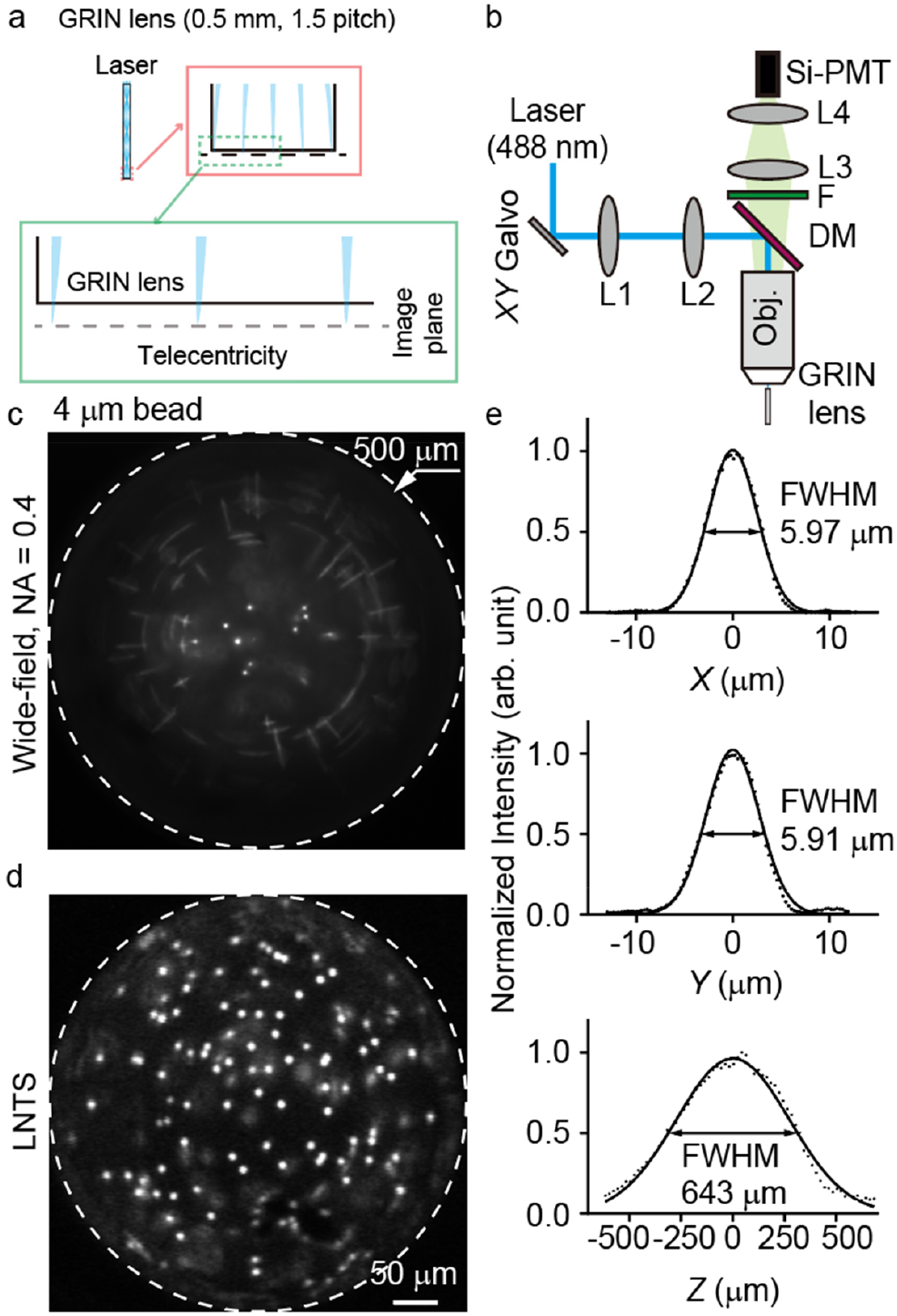
(a) Schematic illustration of the optical path within the GRIN lens. The ray diagram shows telecentricity at the image plane. (b) Optical setup of the LNTS system. Obj, objective lens; L1-4, optical lenses; DM, dichroic mirror; Si-PMT, silicon photomultiplier tube; F, emission filter. (c) Conventional wide-field imaging of fluorescent beads of 4 µm in diameter. (d) LNTS imaging of fluorescent beads of 4 µm in diameter. (e) Point spread function (PSF) quantification. The bead imaging was repeated independently 3 times with similar results.

### Validation through *in vivo* structural imaging

For one-photon imaging in scattering biological tissue, image contrast is a crucial parameter. To evaluate the image contrast over the FOV, we performed *in vivo* imaging of microglia in the brain of *Cx3cr1*-eGFP mice using a GRIN lens of 0.5 mm in diameter and 6.7 mm in length. With the conventional wide-field imaging with an NA of 0.4, the cells were resolvable near the center of the FOV (Fig. 2a). In comparison, with the LNTS approach, the cells were visible even at the edge of the FOV (Fig. 2b). Overall, both methods showed similar contrast in the middle of the FOV, whereas the LNTS approach maintained consistent contrast over the entire 500 µm FOV (Fig. 2c, d), which was unprecedented even with the applications of various correction optical elements. We further explored directly reducing the NA of the camera-based wide-field imaging (Supplementary Fig. 3). As anticipated, low-NA detection further extended the FOV. However, the diminished collection efficiency led to rapid photobleaching within only a few minutes of recording.

**Figure 2.**
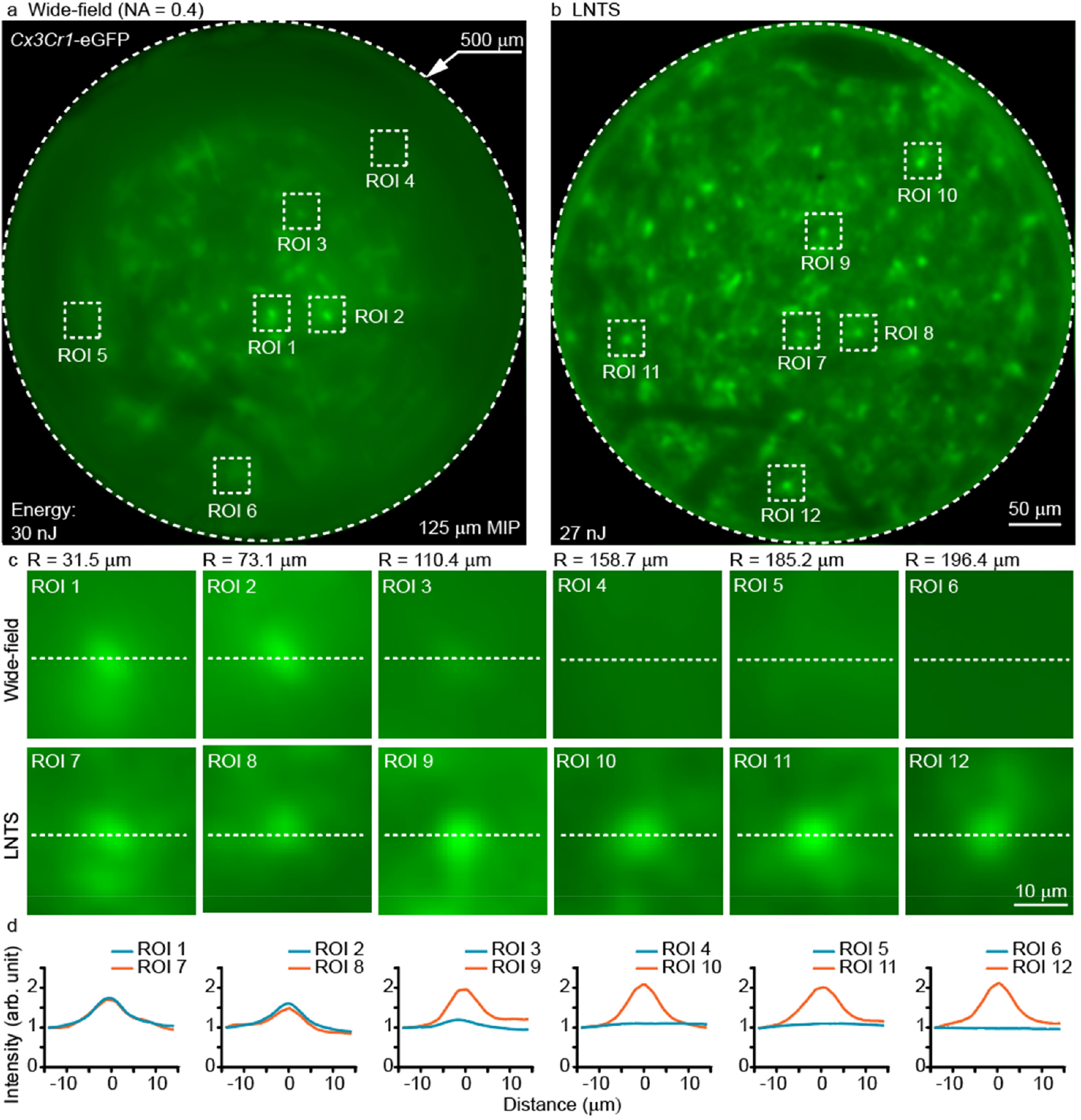
(a) Wide-field image of microglia in the brain of a live *Cx3cr1*-eGFP mouse acquired through a GRIN lens of 0.5 mm in diameter and 6.7 mm in length. The *in vivo Cx3cr1*-eGFP imaging was repeated independently in 2 mice with similar results. (b) LNTS image through the same GRIN lens. (c) Zoomed-in view of the ROIs in a and b. (d) Normalized intensity profiles across the dashed lines in c for image contrast comparison.

### Validation through *in vivo* calcium imaging

To evaluate the performance of *in vivo* calcium imaging, we implanted GRIN lenses of 0.6 mm in diameter and 8.4 mm in length in the thalamus region (AP: -2 mm; ML: ±1.5 mm; DV: -2.85 mm) of the transgenic mice generated by crossing Tbr1-2A-CreER with Ai162-D, and carried out *in vivo* calcium imaging of the neuronal spontaneous activity at a 10 Hz frame rate. Similar to the results of structural imaging, we could consistently image over ∼100% of the GRIN lens diameter. Even cells on the boundaries of the 600 µm diameter were clearly resolved (Fig. 3a). To evaluate the achieved signal quality, we show the raw data of calcium transients from cells across the FOV (Fig. 3b). Even without any signal processing, the raw calcium transients were well above the noise floor. At such high signal levels, we carried out continuous recording for 30 minutes, over which the signal only showed a slight reduction (Fig. 3c), thanks to the LNTS system’s excellent signal collection capability.

**Figure 3.**
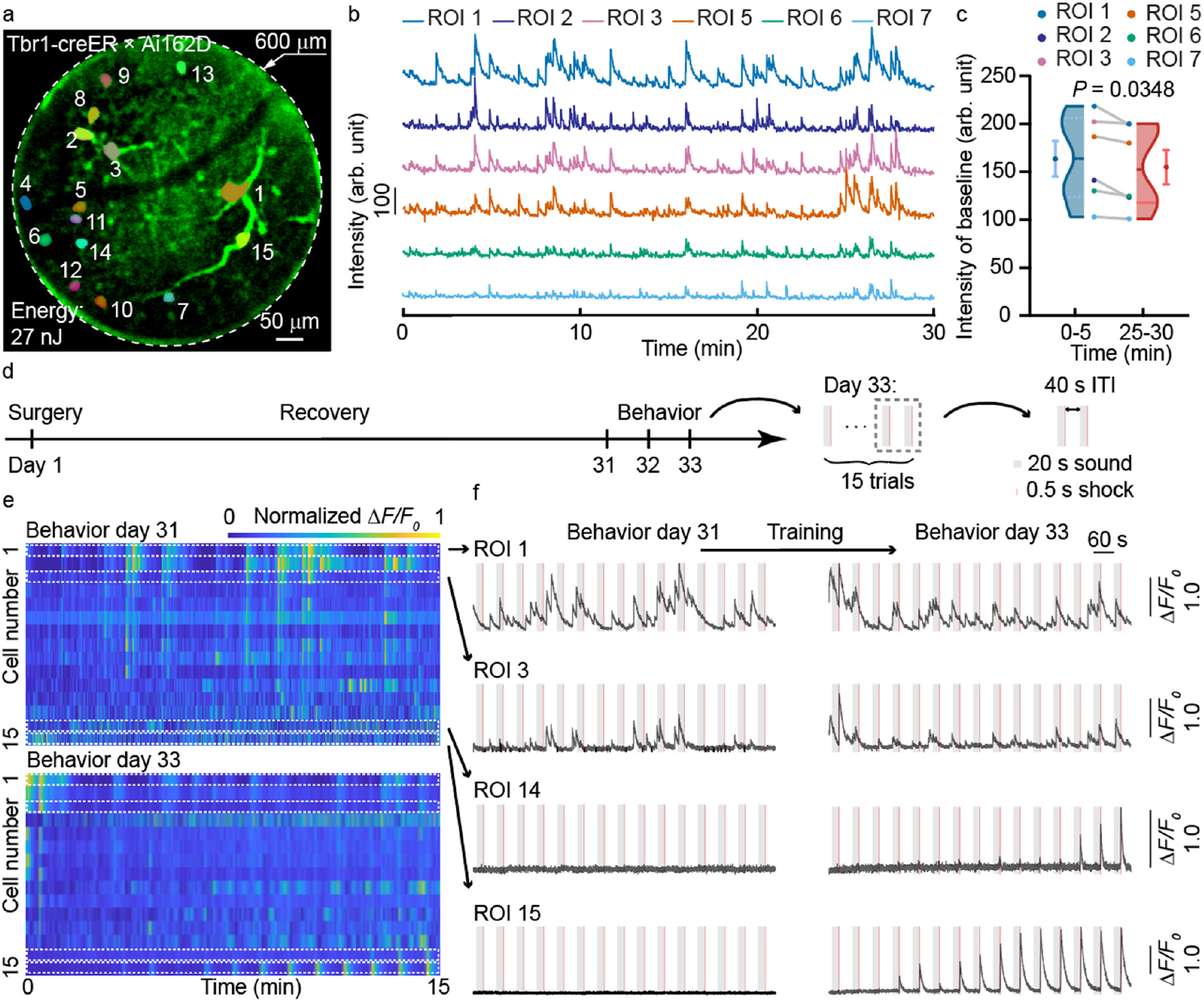
(a) Large FOV calcium image acquired through a GRIN lens of 0.6 mm in diameter and 8.4 mm in length in a Tbr1-creER × Ai162D transgenic mouse. (b) Raw calcium fluorescence traces from six selected ROIs over a 30-minute imaging session. (c) Quantification of fluorescence intensity baseline during early (0–5 min) and late (26–30 min) periods of the session. Each dot and gray line in the center of the chart represents the comparison of the same neuron. The violin plot illustrates the data distribution, with the solid black line indicating the median, and the dashed lines above and below indicating the upper and lower quartiles. The Mean ± SEM is shown next to the violin plot. *P* = 0.0348. Two-tailed paired t-test, t = 2.874, df = 5, n = 6 ROIs from 1 mouse. The *in vivo* photobleaching imaging was repeated independently in 5 mice with similar results. (d) Experimental timeline illustrating the key stages: surgical implantation, recovery, and behavioral testing. ITI: inter-trial interval. (e) Heatmaps of normalized Δ*F/F_0_* traces for all 15 ROIs on Day 31 (pre-training) and Day 33 (post-training). The *in vivo* calcium GCaMP6s imaging was repeated independently in 3 mice with similar results. (f) Calcium traces from four ROIs before and after training.

Next, we used the system for calcium imaging in fear conditioning studies. Specifically, the mouse received an electric shock to its feet after each burst of acoustic cues (Fig. 3d). After three days of training (15 trials per day), individual cells exhibited distinct activity patterns (Fig. 3e). To highlight the excellent signal-to-noise ratio (SNR) of the recorded calcium transients, we showed the raw Δ*F/F_0_* signals without further processing (Fig. 3f). A subset of previously non-responsive cells became synchronized with the stimuli after training, while others remained unsynchronized. It is worth noting that some of the training-responsive cells were located near the outer regions of the FOV (e.g., ROI 14 and 15). Without the LNTS method with ∼100% FOV coverage, they would be imperceptible in the measurements.

To further validate the image fidelity of the calcium recording, we leveraged a new two-photon calcium imaging system recently developed by our lab, specifically for providing ultra-large flat FOV through GRIN lenses, to perform *in vivo* calcium imaging of the same mice (Supplementary Fig. 4). The two-photon system provided better resolution, contrast and imaging depth. For the comparison, we performed a summation of a stack of two-photon images from 0 to 100 µm depth. Although calcium imaging may not be precisely the same depending the neuronal activity during the recording, the two images matched well and the majority of soma can be found on both images.

## Discussion

In this work, we introduce a straightforward solution for maximizing FOV in one-photon calcium imaging through GRIN lenses, which is adaptable to all kinds of GRIN lenses and wavelengths. The low-NA excitation circumvented the native aberration of GRIN lenses, the telecentric scanning avoided vignetting while covering 100% of the GRIN lens facet, and the high-NA signal collection efficiently utilized the fluorescence emission. Collectively, the system enables high-contrast calcium imaging over ∼ 100% of the lens diameter, unprecedented by any other solution. For the common GRIN lenses of 0.5 mm in diameter and ∼7 mm in length, the usable FOV area was increased by ∼400%.

Although a GRIN lens was employed as the implanted deep tissue imaging probe in this study, future work could explore replacing GRIN lenses with high-refractive-index solid glass light pipes for one-photon calcium imaging. The proposed alternative not only supports low-NA excitation light delivery but also provides substantially enhanced fluorescence signal collection efficiency (Supplementary Fig. 5).

The strong correlation between the one-photon and two-photon calcium imaging promises potential multimodal molecular imaging for cell type classification after in vivo calcium recording^11^. Currently, such correlative imaging studies often rely on two-photon *in vivo* calcium imaging, which is expensive, bulky, and difficult to achieve a FOV as large as the GRIN lens facet^30^. In comparison, the LNTS is simple, low-cost (total hardware cost < $10 k), flexible, and can be potentially adapted to freely moving animal studies (e.g., using a fiber bundle to deliver laser scanning and a silicon PMT for signal detection). Thus, LNTS holds great promise to broadly enable such ultra-large FOV multimodal imaging.

Overall, the new LNTS imaging strategy represents a significant advancement in GRIN lens-based deep brain imaging by providing a very simple yet highly effective solution to maximize the FOV without requiring complex adaptive optics or specialized correction lenses customized for specific GRIN lens parameters. With the optical access to the full facet area of the GRIN lens, researchers can now image ∼400% larger neuronal populations with higher experimental success rates. The flexibility and simplicity of this method are expected to accelerate discoveries in the study of deep brain regions.

## Methods

### Animal

The research work complied with all relevant ethical regulations. All procedures involving mice were approved by the Purdue University Animal Care and Use Committee (protocol number: 1506001267). *Cx3cr1*-eGPF, Tbr1-2A-CreER, and Ai162(TIT2L-GC6s-ICL-tTA2)-D mice were obtained from the Jackson Laboratory. The mice were housed in the animal facility of the Bindley Bioscience Center at Purdue University. The facility maintained a light cycle of 12 hours on and off, with lights on from 6 am to 6 pm. The environmental conditions were carefully controlled, with temperatures sustained at 20-25 degrees Celsius. Surgical procedures were performed on adult male and female mice over two months old.

### GRIN lens implantation

In this study, we first created a cranial window with a diameter of 1.5 mm. A wire-cutting tool was then slowly advanced into the brain tissue at a speed of 10 μm/s to reach the target depth. Once positioned, the tool was rotated more than 180 degrees to transect the tissue at the base of the intended GRIN lens implantation site. The tool was then gradually withdrawn at the same speed. Subsequently, a GRIN lens was slowly inserted into the brain, and the implantation site was sealed using an artificial dura (KWIK-SIL, WPI) and dental adhesive. To protect the lens from dust contamination, a plastic cap was placed above the GRIN lens. Following a recovery period of 3–4 weeks, animals proceeded to subsequent experimental procedures.

### Fear conditioning

Mice were head-fixed under the microscope, with their left and right limbs placed separately on two tin foil-covered metal plates that were electrically isolated from each other. Each plate was connected to a distinct output channel of the stimulator. Each stimulation trial lasted 60 seconds, beginning with a 10-second baseline, followed by a 20-second auditory cue (3 kHz tone), and a 30-second post-stimulus baseline. A 24 V foot shock was delivered during the final 0.5 seconds of the tone. Each mouse underwent at least 15 such trials.

## Data analysis

All time-lapse calcium imaging data were processed by the non-rigid registration of the *Suite2p* and *EZcalcium* software to reduce motion artifacts. The calcium images (Fig. 3a, Supplementary Fig. 4a) were processed with image contrast enhancement^35^. Relative fluctuations in fluorescence intensity (Δ*F/F_0_*) were quantified for all regions of interest (ROI). *F*_0_ was defined as the lowest 10% of the fluorescence signal of each ROI. Baselines were quantified separately for the early (0–5 min) and late (25–30 min) epochs (Fig. 3c) to assess potential photobleaching during prolonged imaging. Activity events were defined as upward crossings of Δ*F/F_0_*> 0.4; for each event, the baseline intensity was calculated as the mean intensity over the 2 s preceding onset. To avoid elevated baselines caused by incomplete recovery from preceding events, only events with pre-onset Δ*F/F_0_*< 0.2 were included in the analysis. The intensity scale of the LNTS images (Fig. 2c) was normalized to that of the wide-field images, using the baseline intensity at the left side as the reference. For comparison across ROIs, all line intensity profiles (Fig. 2d) were normalized such that the baseline intensity at the left side of each profile was unity. The energy values shown in the main and supplementary figures represent the excitation energy delivered over the entire GRIN lens facet to have ∼ 1,000 detected photons per bead/cell.

## Statistical Analysis

All data in this study were tested for normal distribution. Data that passed the normality test were analyzed for significance using the two-tailed paired t-test. All statistical analyses were performed with *GraphPad Prism 10*. No results from successful image acquisitions and measurements were excluded or filtered. Specific values for *P*, n, etc., are presented in the figure captions.

## Acknowledgment

This work was funded by NIH (U01NS126054, U01NS118302). The funders had no role in study design, data collection and analysis, decision to publish, or manuscript preparation. M.C. thanks the Howard Hughes Medical Institute and Purdue University for scientific instruments.

## Author Contributions Statement

M.C. invented the imaging method and designed the imaging system. C.W. implemented the imaging system and carried out the imaging experiments. Z.C. performed surgeries on mouse brains. C.W. and Z.C. collaborated on the experiment, data analysis, and figure preparation. Y.L. facilitated the *in vivo* imaging performance quantification and the comparison between one-photon and two-photon images. J.L. facilitated the optical detection system. M.C. wrote the manuscript with input from all authors.

## Supplementary Material for

**This file includes Supplementary Figures 1-5.**

**Supplementary Figure 1.**
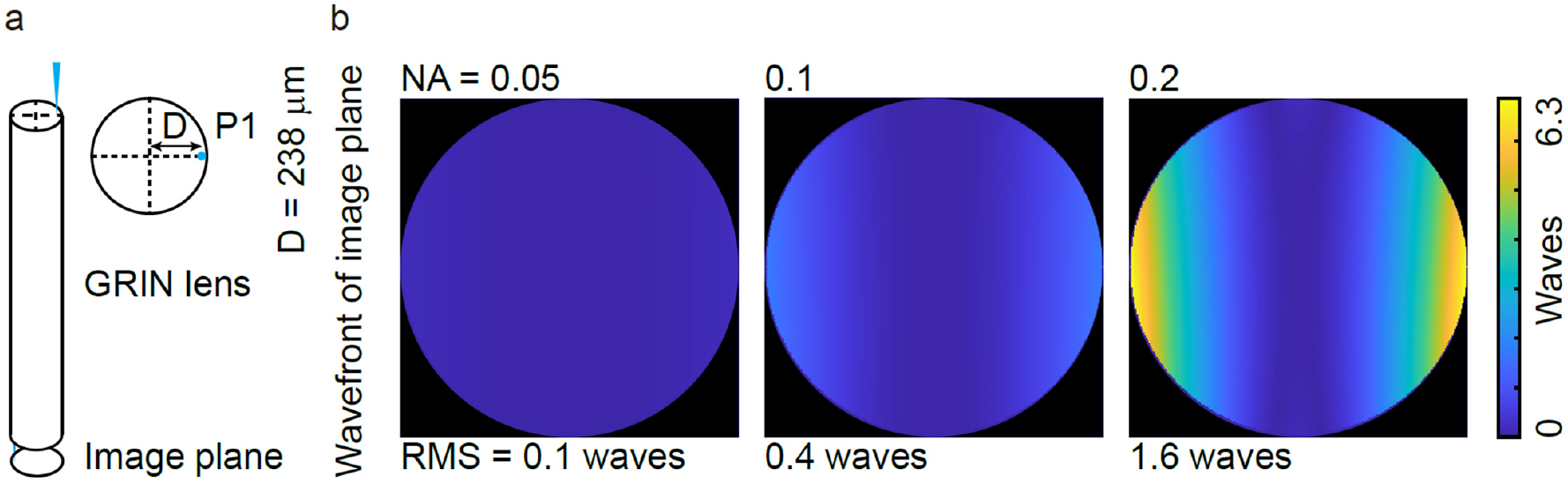
Simulated wavefront through a GRIN lens of 0.5 mm in diameter 6.7 mm in length. (a) Beam path configuration. The exit location is 238 µm from the center of the FOV (b) GRIN lens induced aberration as a function of input NA.

**Supplementary Figure 2.**
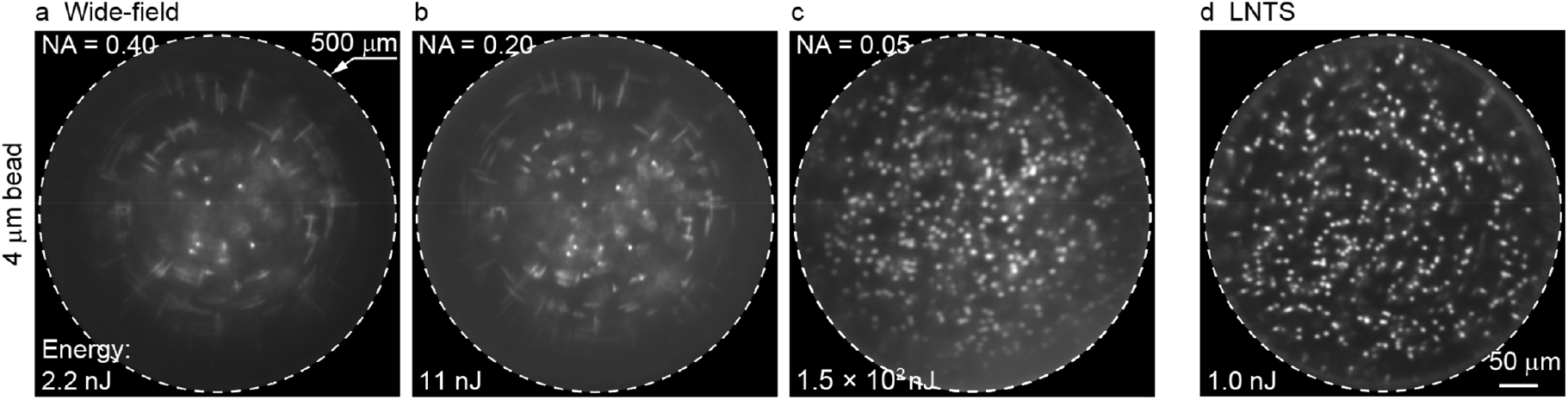
Comparison of wide-field imaging and the LNTS method for imaging through a GRIN lens of 0.5 mm in diameter 6.7 mm in length. (a-c) Wide-field imaging with a detection NA of 0.4, 0.2 and 0.05, respectively. (d) LNTS based imaging of the same sample. The illumination energy over the 500 µm FOV required for the beads in the middle to have ∼1,000 emission photons collected is shown for each image.

**Supplementary Figure 3.**
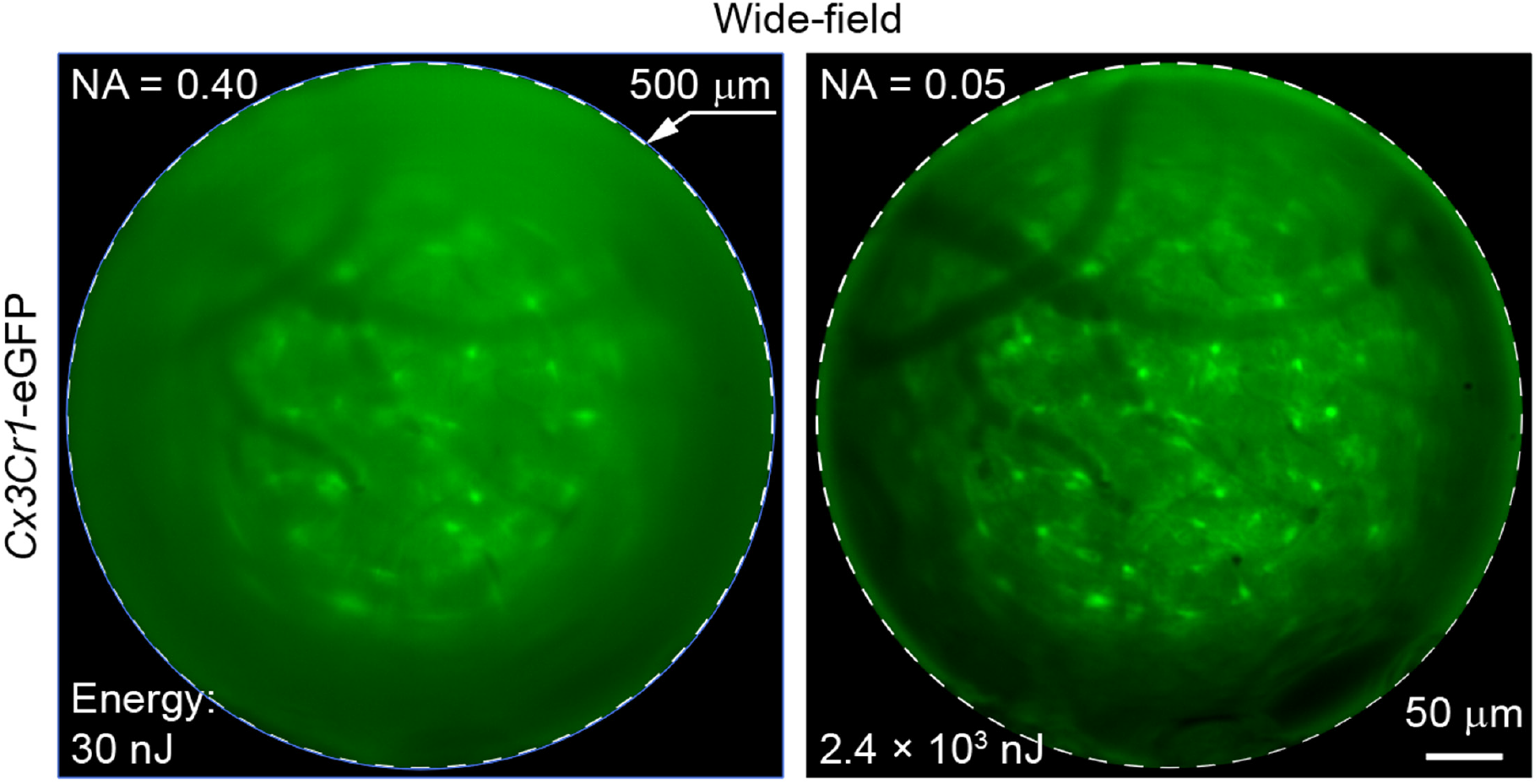
*In vivo* wide-field imaging of microglia through a GRIN lens of 0.5 mm in diameter 6.7 mm in length. Images of microglia through the same GRIN lens at a detection NA of 0.4 and 0.05, respectively.

**Supplementary Figure 4.**
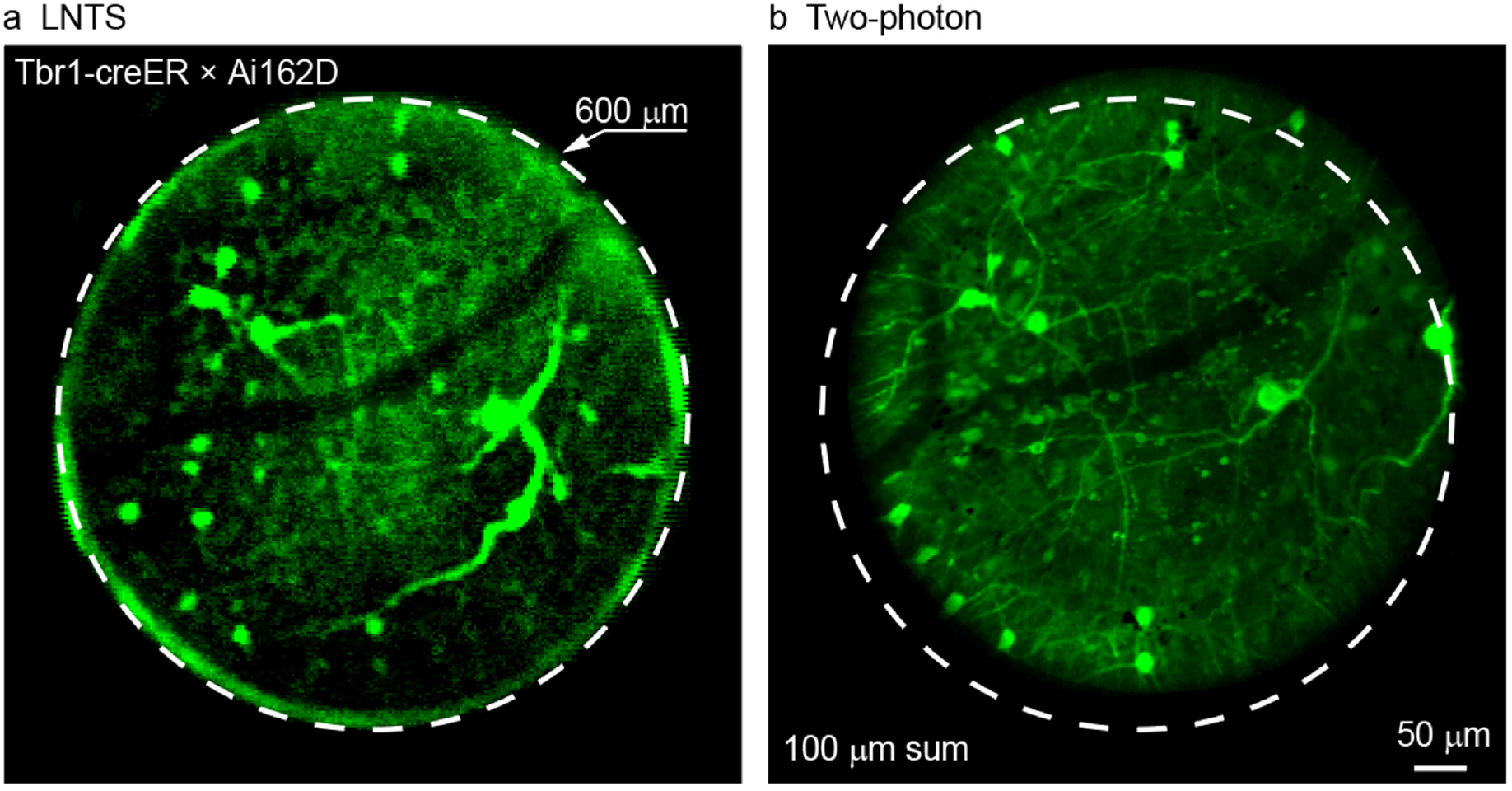
*In vivo* comparison of LNTS and ultra-large flat FOV two-photon system for imaging neurons through a GRIN lens of 0.6 mm in diameter 8.4 mm in length. (a, b) Images of neurons acquired by LNTS and by ultra-large flat FOV two-photon system, respectively.

**Supplementary Figure 5.**
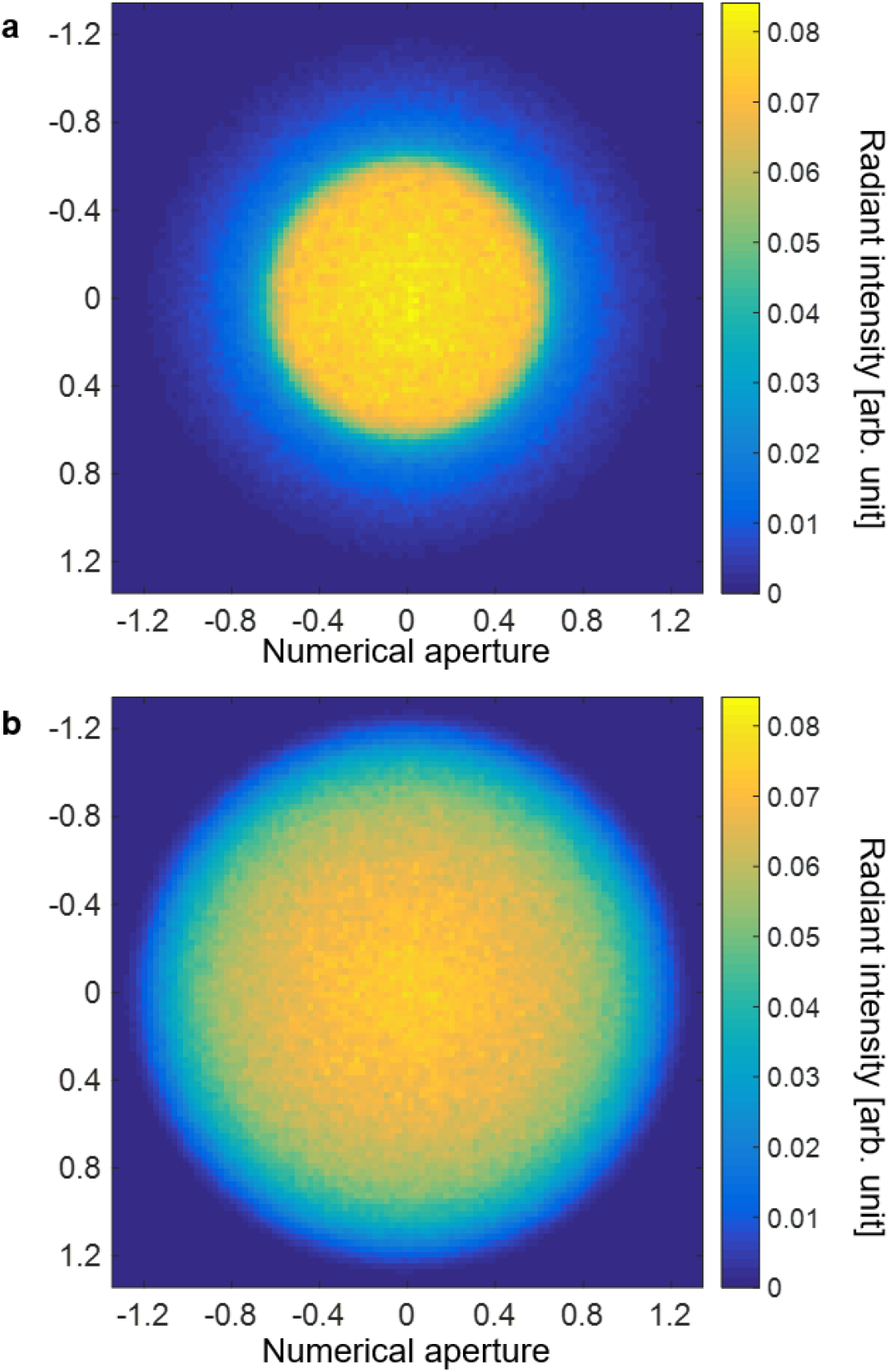
Non-sequential ray tracing through a GRIN lens of 0.5 mm in diameter 6.7 mm in length and a uniform H-ZLAF78B glass light pipe of identical dimensions. (a) The radiant intensity distribution of the light collected by the GRIN lens. The signal source is a uniform cylinder of 0.5 mm in diameter and 0.1 mm in length located right under the GRIN lens facet. Among the isotropic emissions, 15.7% is captured through the GRIN lens. (b) The radiant intensity distribution of the light collected by the H-ZLAF78B glass light pipe under the same condition. The collection efficiency is 27.6%.

